# Cancer progression models and fitness landscapes: a many-to-many relationship

**DOI:** 10.1101/141465

**Authors:** Ramon Diaz-Uriarte

## Abstract

The identification of constraints, due to gene interactions, in the order of accumulation of mutations during cancer progression can allow us to single out therapeutic targets. Cancer progression models (CPMs) use genotype frequency data from cross-sectional samples to try to identify these constraints, and return Directed Acyclic Graphs (DAGs) of genes. On the other hand, fitness landscapes, which map genotypes to fitness, contain all possible paths of tumor progression. Thus, we expect a correspondence between DAGs from CPMs and the fitness landscapes where evolution happened. But many fitness landscapes —e.g., those with reciprocal sign epistasis— cannot be represented by CPMs. Using simulated data under 500 fitness landscapes, I show that CPMs’ performance (prediction of genotypes that can exist) degrades with reciprocal sign epistasis. There is large variability in the DAGs inferred from each landscape, which is also affected by mutation rate, detection regime, and fitness landscape features, in ways that depend on CPM method. And the same DAG is often observed in very different landscapes, which differ in more than 50% of their accessible genotypes. Using a pancreatic data set, I show that this many-to-many relationship affects the analysis of empirical data. Fitness landscapes that are widely different from each other can, when evolutionary processes run repeatedly on them, both produce data similar to the empirically observed one, and lead to DAGs that are very different among themselves. Because reciprocal sign epistasis can be common in cancer, these results question the use and interpretation of CPMs.

## 1 Introduction

Epistatic interactions between genetic alterations constraint the order of accumulation of mutations during cancer progression (e.g., in colorectal cancer APC mutations precede KRAS mutations [18]). Finding these constraints can single out therapeutic targets and disease markers and has lead to the development of cancer progression models (CPMs) [3], such as CBN [21, 22] or CAPRI [9, 40], that try to identify these constraints using genotype frequency data using cross-sectional samples. CPMs return directed acyclic graphs (DAG) of genes where arrows indicate dependencies or constraints (Figure 1). The DAGs specify that only genotypes that fulfil certain restrictions can exist (Figure 1b and 1f).

**Figure 1:**
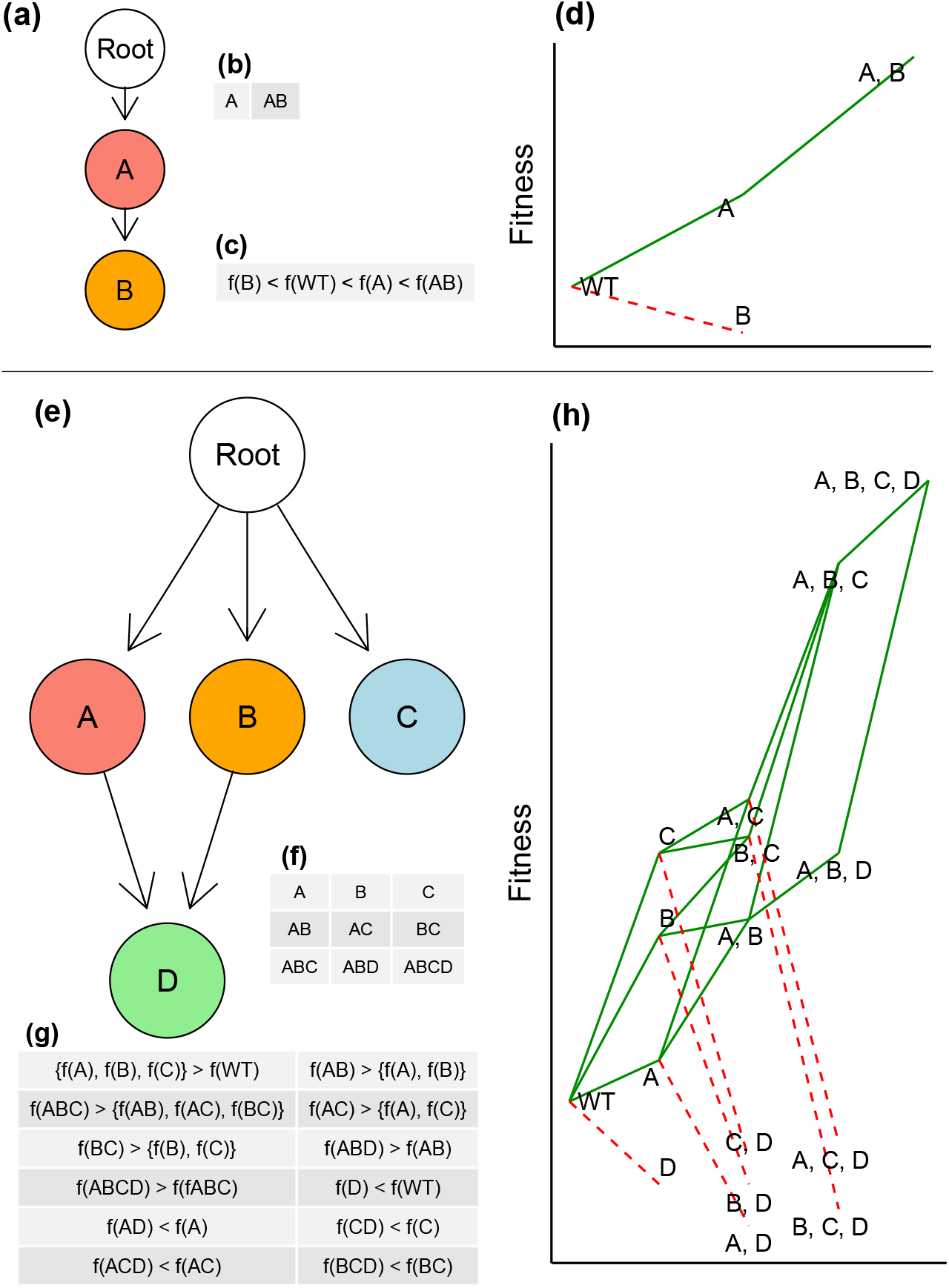
DAGs of genes from cancer progression models and representable fitness landscapes. (a) An arrow (directed edge) from gene *i* to gene *j* means a mutation in *j* cannot be observed unless i is mutated. Here a mutation in gene B can only be observed if A is mutated (i.e., B increases fitness only if A is mutated). Tumor evolution starts at “Root” and genes whose parent is Root do not depend on mutations of other genes. (b) The genotypes that can exist according to that DAG, where the notation “A” means a genotype with gene “A” (but not B) mutated and “AB” a genotype with both genes A and B mutated. (c) The fitness relationships between genotypes implied by the DAG, with “f(A)” denoting fitness of a genotype with gene A mutated (and “WT” the wild-type, non-mutated genotype). (d) A fitness landscape that can be represented by the DAG in (a); genotypes as in (a) and (b) (this fitness landscape representation is based on [8]). Green segments connect genotypes along accessible paths and red dotted segments denote decreases in fitness. I will use “accessible genotypes” to refer to genotypes along accessible paths; thus, genotypes reachable through green segments are accessible and those only reachable through red segments are non-accessible. (e) to (h) as in (a) to (d) but with four genes. Note the use of the logical AND for the arrows on D: both A and B must be present for D to happen. In (g) only some of the implied relationships are shown. That the fitness landscape be representable by the DAG depends only on the fitness ranks of genotypes [13, 20], not on the specific values of their fitness: note the agreement between (c) and the ranks of genotypes’ fitness from (d), and between (g) and the ranks of genotypes’ fitness from (h).

Whereas DAGs of genes from CPMs do not contain information about the fitness of individual genotypes, fitness landscapes (or genotype-fitness maps) associate to each genotype its fitness value [14]. Thus, similarly to DAGs of genes, a fitness landscape, if we assume that populations will only move uphill in fitness, specifies what genotypes can be observed [13, 20]. Crosssectional samples taken during tumor progression should contain only genotypes that are part of accessible paths (accessible path: a trajectory through a collection of genotypes, where each genotype is separated from the preceding genotype by one mutation, along which fitness increases monotonically [20]). If we had detailed knowledge about the fitness landscape, we could predict the possible paths of tumor progression and identify genes that would block those paths [23, 29]. However, obtaining a complete picture of the cancer fitness landscape is not currently possible [29, 42]. Here CPMs can offer a feasible alternative: a model that identifies the key constraints in the order of accumulation of mutations might be enough to capture the possible tumor progression paths [2] and the genotypes that can and cannot exist.

But then, we should expect a correspondence between DAGs from CPMs and the fitness landscapes where tumor evolution took place. DAGs should provide accurate predictions about what genotypes can and cannot exist during tumor progression, and the same DAG should not be inferred from very different fitness landscapes. Are these expectations justified? Do they hold with empirical data? And what are the consequences of using CPMs when the expectations do not hold?

With some fitness landscapes, that correspondence might hold (if sample sizes are sufficiently large). For example, we will say that the landscapes in Figures 1d and Figure 1h are representable by the DAGs in Figure 1a and Figure 1e, respectively: the DAGs and the landscapes make the same predictions about what genotypes we should observe. The landscapes are representable because the DAGs capture the epistatic interactions that determine what genotypes are accessible. In particular, the constraints reflected in the DAGs imply sign epistasis [48], an interaction between genes where a mutation is beneficial or deleterious (i.e., can have different sign) depending on the genetic background or what other genes are mutated (this is the basis of the phenomenon of “oncogene addiction” [42]). Figure 1a says that a mutation in B increases the fitness of a cell if A is already mutated but is detrimental otherwise; Figure 1e says that a mutation in D increases fitness if A and B are mutated but is detrimental otherwise (note that in the landscape the only accessible genotypes with D mutated also have A and B mutated).

For other fitness landscapes, however, the correspondence cannot hold. Although DAGs represent sign epistasis, they cannot represent reciprocal sign epistasis, a genetic interaction where two mutations that individually increase fitness reduce it when combined [13, 38, 39]. CPMs assume that acquiring a mutation in one gene, say A in Figure 1e, does not decrease the probability of acquiring a mutation in another gene, say C [34]. The DAGs can only say what mutations need to be present before another mutation is viable. Thus, neither the DAG in Figure 1e, nor any other DAG of genes, could represent the fitness landscape that would result if reciprocal sign epistasis [13, 38] between genes A and C turned genotype AC into a low fitness or non-viable genotype (and similarly for reciprocal sign epistasis between B and C). (See Supplementary Material, “Representable landscapes with reciprocal sign epistasis?” for apparent exceptions).

This is a potentially serious limitation of CPMs because reciprocal sign epistasis is probably common in cancer [11], given the extent of synthetic lethality both in the human genome [6] and in cancer cells [4, 28, 44] (synthetic lethality is an epistatic interaction where the combination of two mutations is lethal when each individual mutation is not —synthetic lethality between mutations that individually increase fitness constitutes reciprocal sign epistasis). Moreover, reciprocal sign epistasis is a key structural feature of fitness landscapes: it can lead to multiple peaks and affects ruggedness and predictability of the evolutionary process [13, 14, 19, 38], which itself affects our opportunities to block tumor progression [23, 29]. But the assessment of CPMs has used data simulated from generative models that are encoded by DAGs [21, 24, 40, 46], therefore assuming very restricted fitness landscapes. Two exceptions, none of which considered reciprocal sign epistasis explicitly, are [43], who conducted simulations using agent-based models with parameters tuned for colorectal cancer, and [16], where the restrictions encoded in DAGs were embedded within evolutionary models that allowed to relax some of the constraints of the DAGs. Using fitness landscapes, as done in this paper, is a more direct route to examine the consequences of more complex evolutionary scenarios for CPMs and it avoids the shortcomings of both agent-based models [43], where it can be hard to understand the model in terms of generalizable features, and of [16], which could only use a limited subset of deviations from DAGs.

The main goal of this paper is to understand the relationship between fitness landscapes and CPMs inferred from evolutionary process on those fitness landscapes. I used evolutionary simulations on 500 fitness landscapes that include from none to extensive reciprocal sign epistasis, varying also mutation rate and time to tumor detection (as they condition what genotypes are observable from a landscape). Because the focus of this paper is not method comparison *per se*, but to identify the limits that complex fitness landscape can impose on the use of CPMs, I used large sample sizes of N = 1,000 genotypes and inferred CPMs with CBN [21, 22] and CAPRI [40], the two widely available state-of-the-art methods that accommodate multiple parents in DAGs. I evaluate the effects of fitness landscape, mutation rate, tumor detection, and reciprocal sign epistasis on the quality and variability of inferred DAGs. As expected, I find that DAGs often mispredict the genotypes that can exist under a fitness landscape with reciprocal sign epistasis. Unexpectedly, however, I find a many-to-many mapping between fitness landscapes and DAGs that affects both representable and non-representable landscapes. I then examine the practical consequences of this many-to-many mapping using a pancreatic cancer data set: I find that fitness landscapes that are widely different from each other can produce genotype frequencies similar to the empirically observed one, and also lead to very different DAGs when evolutionary processes run repeatedly on them. These results cast doubts on whether restrictions from DAGs can be used to capture the true restrictions in fitness landscapes.

### 1.1 Assumptions

Several assumptions were used in the arguments above and are taken for granted in the rest of this paper. As is customary in the CPM literature [3, 13, 34, 40] in this paper I will not deal with possible complications arising from polyclonality [10, 36] and non-cell autonomous interactions and I assume that the DAGs and fitness landscapes represent epistatic interactions. Also in agreement with common models in this field [1, 7, 33] back mutations are not allowed. Neither are multiple simultaneous mutations allowed: crossing valleys in the fitness landscape in a single multi-mutation step is not possible (this assumption might turn out to be inappropriate if phenomena such as chromothripsis [45] are common), but we do not need to exclude clonal interference [14, 41] (see also Supplementary Material “Paths through non-accessible genotypes”). Finally, I assume no observational errors and I assume that we know which are the driver genes (so we do not need to remove passengers before CPM fitting [12, 16]). Addressing these issues is beyond the scope of this paper.

## 2 Results

### 2.1 Fitness landscapes and simulation framework

To understand the consequences of changes in fitness landscape characteristics, tumor detection, and mutation rates, on the quality and variability of DAGs, I generated 500 fitness landscapes (see Methods). Of these, 100 are directly derived from DAGs of restrictions and, thus, are perfectly representable and contain no reciprocal sign epistasis. Another 200 were obtained from representable fitness landscapes in which randomly selected genotypes predicted by the DAG were made inaccessible: by introducing synthetic lethals I created DAG-derived, but not representable, fitness landscapes with varying amounts of reciprocal sign epistasis (see Supplementary Material “Fitness landscapes characteristics”; see also Supplementary Material, “Canonical DAG” for checking if a landscape is representable). Another set of 200 landscapes were obtained from Rough Mount Fuji (RMF) models, which have been useful to model empirical fitness landscapes [14, 35] and have very different characteristics from DAG-derived fitness landscapes (see Supplementary Material “Fitness landscapes characteristics”); all of the RMF models were not representable. RMF landscapes had both more reciprocal sign epistasis and larger numbers of peaks than DAG-derived fitness landscapes (average fraction of pairs of loci with reciprocal sign epistasis [19]: 0.27 vs 0.02 for RMF- and non-representable DAG-derived landscapes; two-sample t-test, *t*_240_ = 45, p < 0.0001; number of peaks among the accessible genotypes: 13.4 and 3.13 in the RMF and DAG-derived, respectively; two-sample t-test, *t*_310_ = 37, *p* < 0.0001).

Next, I simulated evolutionary processes on the 500 fitness landscapes using a logistic-like model, following [33], where death rate depends on total population size. I used a factorial design where on each one of the 6,000 combinations of 500 landscapes by three mutation rates by two tumor detection schemes (see Methods), I run 20,000 evolutionary processes that resulted in 20,000 simulated sampled genotypes per condition. The 20,000 simulated genotypes were split into 20 sets of 1,000 genotypes and each one of the 20 sets was analyzed with CBN and CAPRI (yielding, therefore, 20 DAGs per method for each one of the 6,000 combinations of landscape*mutation*detection). (Plots of the 500 fitness landscapes and the modal inferred DAG are provided in the Supplementary Material, “Plots of fitness landscapes and inferred DAGs”.)

### 2.2 Genotype mispredictions in inferred DAGs

For each genotype, I compared the status accesible/non-accessible predicted by each DAG with the true status from the fitness landscape. The proportion of false discoveries (PFD) is the number of genotypes that are erroneously predicted to exist relative to the number of genotypes predicted to exist by a DAG. The proportion of negative discoveries (PND) is the number of accessible genotypes that a DAG fails to predict relative to the total number of accessible genotypes (the PND measure corrects for genotypes that are not observable for a given combination of detection and mutation rates —see Methods). Figure 2a, b shows the PFD and PND for each method and landscape type. There was strong evidence of differences between landscapes and method for both measures (interaction method by landscape from linear mixed-effects models of PFD and PND as dependent variables with method and landscape type as explanatory variables: Type II Wald F tests with Kenward-Roger d.f. adjustment: *F*_2,5493_ of 189.2, 524.20, for PFD and PND, respectively, both *p* < 0.0001).

**Figure 2:**
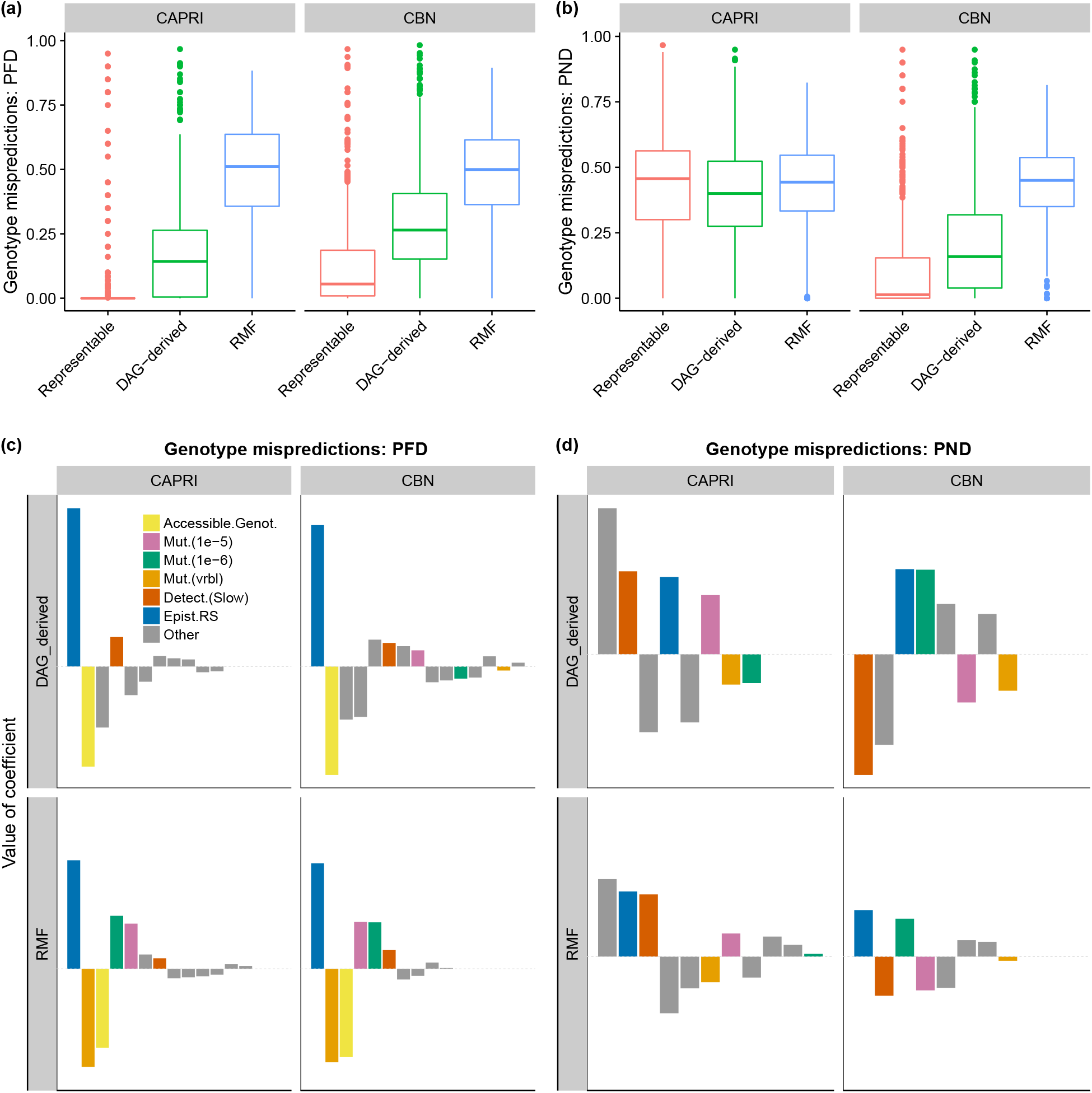
Genotype mispredictions: proportion of false discoveries (PFD) and proportion of negative discoveries (PND). PFD: number of genotypes that are erroneously predicted to exist relative to the number of genotypes predicted to exist by a DAG. PND: number of accessible genotypes that a DAG fails to predict relative to the total number of accessible genotypes. (a) and (b) Boxplots of PFD and PND by method and type of fitness landscape. (c) and (d) coefficients from linear mixed-effects models, with separate models fitted for each combination of method and type of fitness landscape. Within each panel coefficients have been ordered from left to right according to decreasing absolute value of coefficient. The dotted horizontal gray line indicates 0 (i.e., no effect). Scale of the y-axes is the same for the four panels of a response variable. Continuous regressors (fraction of epistasis and accessible genotypes) centered and scaled (mean 0, variance 1). Only coefficients that correspond to a term with a *p* – value < 0.05 in Type II F (ANOVA) tests are shown. Coefficients involving Landscape type “DAG-derived” and Detection Fast are not shown (they are minus the corresponding coefficient shown for the other value of the factor —see Methods). Coefficients that correspond to main effects have been color coded as shown in the legend; the rest of the coefficients (“Other”) correspond to interaction terms; “Epist.RS”: fraction of pairs with reciprocal sign epistasis; “Mut.(vrbl)”: variable or gene-specific mutation.

I examined how PFD and PND were affected by mutation rate, detection regime, number of accessible genotypes, and reciprocal sign epistasis and their two-way interactions. Figure 2 shows the relative magnitude of the coefficients in the models (see Supplementary Material, “Coefficients of linear models” for the complete set of coefficients). Mainly for PND, the effects of some predictors differed between CBN and CAPRI: models that examined both methods simultaneously showed highly significant interactions between method and the effects of mutation and detection (*p* < 0.0001 from Type II Wald F tests with Kenward-Roger d.f. adjustment), in some cases with reversal of the effects. For PND there were also other large interactions that involved mutation and detection regimes. Increasing reciprocal sign epistasis was associated with increasing PFD and PND for both methods and landscape types. Remarkably, there was no significant interaction between method and reciprocal sign epistasis (*p* > 0.15): we cannot reject the hypothesis that the increase in mispredictions with reciprocal sign epistasis is similar in CBN and CAPRI. But there were strong interactions between type of landscape and reciprocal sign epistasis (*p* < 0.0001): both PFD and PND showed a faster increase with reciprocal sign epistasis in DAG-derived than in RMF landscapes.

### 2.3 The many-to-many mapping between fitness landscapes and DAGs

To examine within-landscape DAG variability I computed the average relative pairwise DAG distance between all possible DAGs of: a) the 20 replicate inferences for each landscape by mutation by detection regime —i.e., same mutation and detection (“Same” in Figure 3); b) the 20 * 6 replicates for each landscape over the six conditions of mutation by detection regime (“Over” in Figure 3). Figure 3a, b shows the probability of obtaining different DAGs in two runs and the relative number of edges on which they differ, respectively. Multiple, substantially different, DAGs were inferred from the same landscape, even under the same mutation rates and detection regimes. (As before, there was strong evidence of interactions between method and landscape type, *p* < 0.0001, in relative pairwise distance in both “Same” and “Over”). Linear models (Figure 3c, d) showed that increasing the number of accessible genotypes increased the variability of inferred DAGs. But, as we saw before, the relevance and sign of other terms depended on the method: mutation, detection, and landscape type showed significant interactions with method (*p* < 0.0001 for all two-way interactions with method; *p* = 0.0009 for the interaction mutation by landscape type by method; and *p* < 0.0001 for the interaction mutation by detection by method). Reciprocal sign epistasis only had a relevant effect when using CBN under the DAG-derived landscape (there was a three-way interaction method by landscape type by epistasis; *p* = 0.002 and *p* = 0.033 for “Same” and “Over”, respectively).

**Figure 3:**
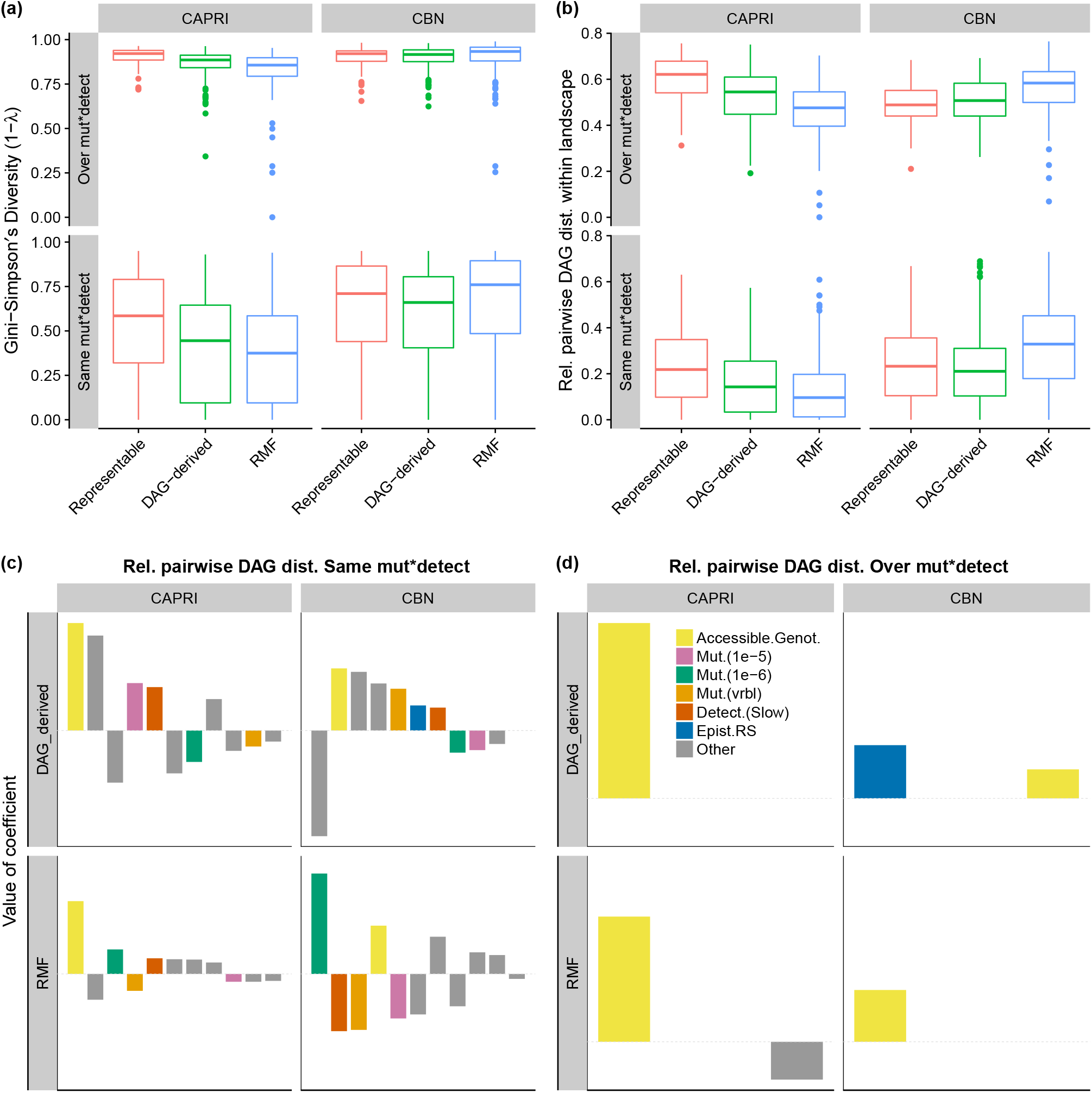
Variability of DAGs inferred from a landscape. Condition “Same mut*detect” corresponds to the values from the 20 replicates under each combination of landscape by mutation rate by detection. Condition “Over mut*detect” corresponds to the 20*6 replicates of a landscape over the 3×2 mutation by detection regime. (a) Gini-Simpson’s diversity index, the probability that two DAGs from the give scenario are different. (b) Relative pairwise DAG distance (see methods for definition). The value shown for a landscape is the average relative pairwise DAG distance for all pairs of inferred DAGs for a landscape. (c) and (d): Coefficients from linear models; see legend of Figure 2. In panel (d) only accessible genotypes, reciprocal sign epistasis, and their interaction could, by design, be examined.

I next examined whether the same DAG was inferred from distinct fitness landscapes. I regarded as distinct landscapes those fitness landscapes that differed in > 50% of their accessible genotypes (i.e., landscapes with a relative pairwise difference of accesible genotypes > 0.5 —see Methods). I first counted for each DAG in how many pairs of distinct landscapes it had been inferred. Of the inferred DAGs, 8% and 11% for CBN and CAPRI, respectively, were present in one or more pairs of distinct landscapes and some DAGs were present in hundreds of different pairs of distinct landscapes (Figure 4a).

**Figure 4:**
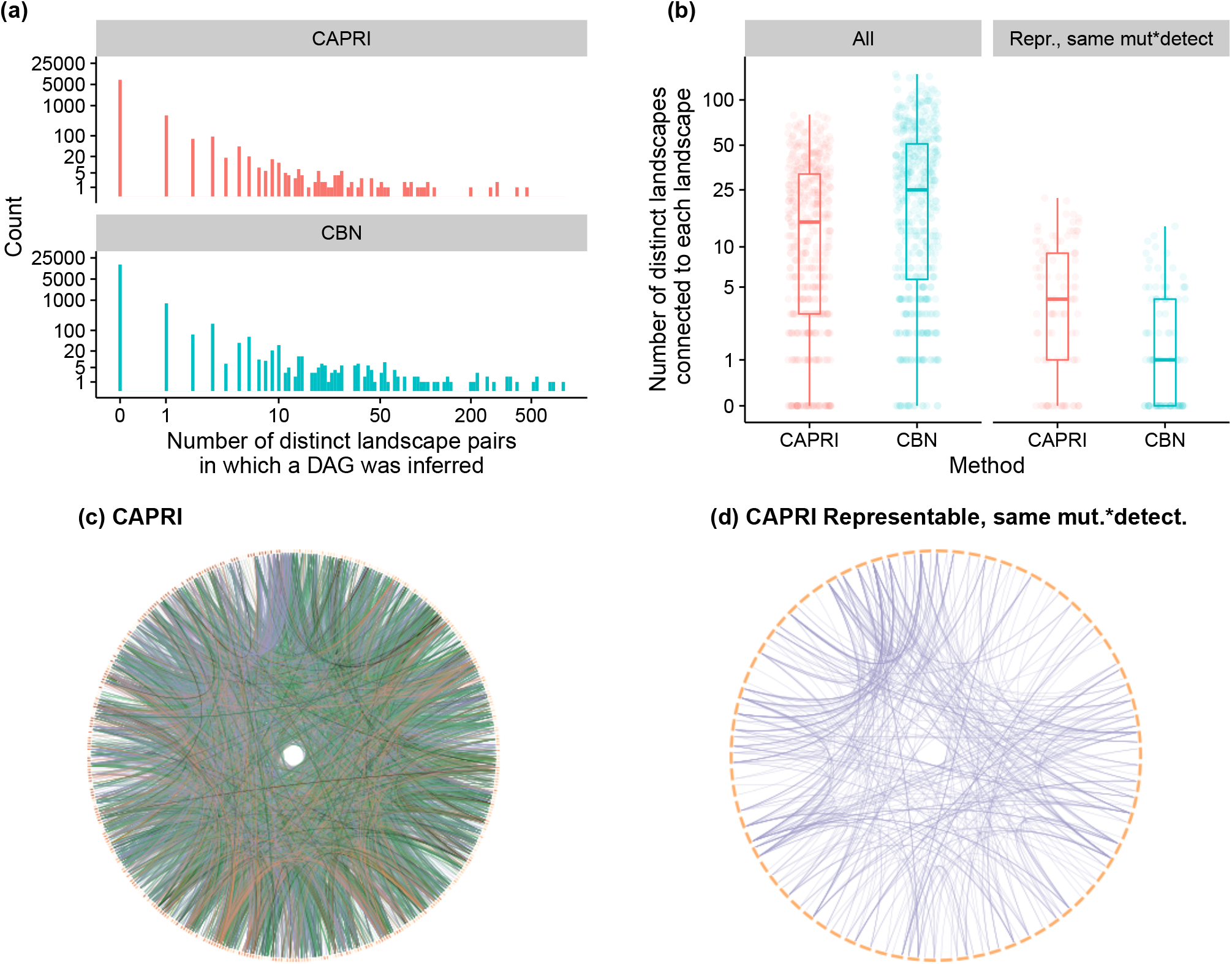
Distinct fitness landscapes with the same inferred DAG. (a) Histograms of the number of distinct landscapes in which DAGs were found, with both y-and x-axes in log-scale; for example, for both CAPRI and CBN about 1,000 different DAGs were found in one pair of distinct landscapes, and for CBN 50 DAGs were found in 10 distinct lanscape pairs. (b) Box-plots with overimposed, horizontally-jittered, data of number of distinct landscapes connected to each landscape for all landscapes under all conditions (left) and only representable landscapes connected under the same mutation and detection conditions (right). (c) Circos [27] plot of connections between distinct landscapes for CAPRI; 448 landscapes (ideograms) are shown, with landscapes ordered by increasing number of accessible genotypes starting from the 12 o’clock position; green links are connections where the same DAG was obtained under the same detection regime but different mutation rates; orange links for different detection regime and same mutation rates; black links for different detection and mutation; purple links for identical detection and mutation. (d) Like (c), but only for representable landscapes and same mutation and detection conditions; 78 landscapes shown.

I also counted, for each fitness landscape, with how many distinct landscapes it was connected, where two landscapes were connected if the same DAG had been inferred in both landscapes in at least one occasion. The vast majority of landscapes (96% and 90% for CBN and CAPRI, respectively) were connected with at least one distinct landscape (Figure 4b). This lead to a complex pattern of relationships between fitness landscapes via shared DAGs (Figure 4c; see Supplementary Material, “Circos plots of landscape connections” for CBN’s results). These results were not the consequence of the same DAG appearing in different landscapes under different mutation and detection regimes. Restricting the analysis to the same DAG appearing under identical mutation and detection conditions, there were 426 and 451 landscapes connected to a distinct landscape in CAPRI and CBN, respectively. These patterns were also observed when the analysis was restricted to DAG-representable landscapes under the same mutation and detection conditions (Figure 4b, right subpanel; Figure 4d): 78 and 55, for CAPRI and CBN, respectively, of the 100 representable landscapes were connected to at least one other distinct representable landscape under the same mutation and detection conditions. Hence, the data show a “one DAG, many landscapes” phenomenon. Because we see both a “one DAG, many landscapes” phenomenon, and large variability in inferred DAGs for the same landscape, I will use the expression “many-to-many” to refer to the DAG-landscape relationship.

### 2.4 The many-to-many problem in a pancreatic cancer data set

I will show the implications of the many-to-many phenomenon for the analysis of empirical data with the 90-patient pancreatic data originally from [26] that was analyzed by [22] using the seven most frequent genes. Since we do not know the true fitness landscape that generated those data we cannot assess the quality of the inferences. What I have done instead is find a set of 162 random fitness landscapes (plus mutation rates and detection regimes) that can produce genotype frequencies similar to the reference one (the original data). A genotype frequency from a simulation was considered similar to the reference one if a *χ*^2^ test of the null hypothesis that both data sets had the same genotype frequencies had a p-value larger than 0.6 and all seven genes had been observed. Next, from each one of those fitness landscapes I generated 20,000 genotypes, and analyzed 20 subsets of 90 genotypes and 20 subsets of 1000 genotypes with CAPRI and CBN. The 162 landscapes and the modal DAGs inferred with CBN and CAPRI for each of the sample sizes are provided in the supplementary material (“Pancreas: landscapes and inferred DAGs”). I am not claiming any of these 162 landscapes is the true pancreatic cancer fitness landscape: I am using them to examine the implications of the many-to-many problem for the analysis of empirical data.

The characteristics of the 162 fitness landscapes are summarized in Figure 5a,b. There was substantial reciprocal sign epistasis (median 0.182, IQR: 0.165) and the landscapes were multi-peaked (median number of peaks: 7, IQR: 5.25). Fitness landscapes were widely different among themselves (Figure 5c,d), with the median pairwise difference in accessible genotypes of 32 genotypes (IQR: 9.6) which, relative to the number of distinct accessible genotypes, was a median pairwise difference of 56% (IQR 34%) of the accessible genotypes.

**Figure 5:**
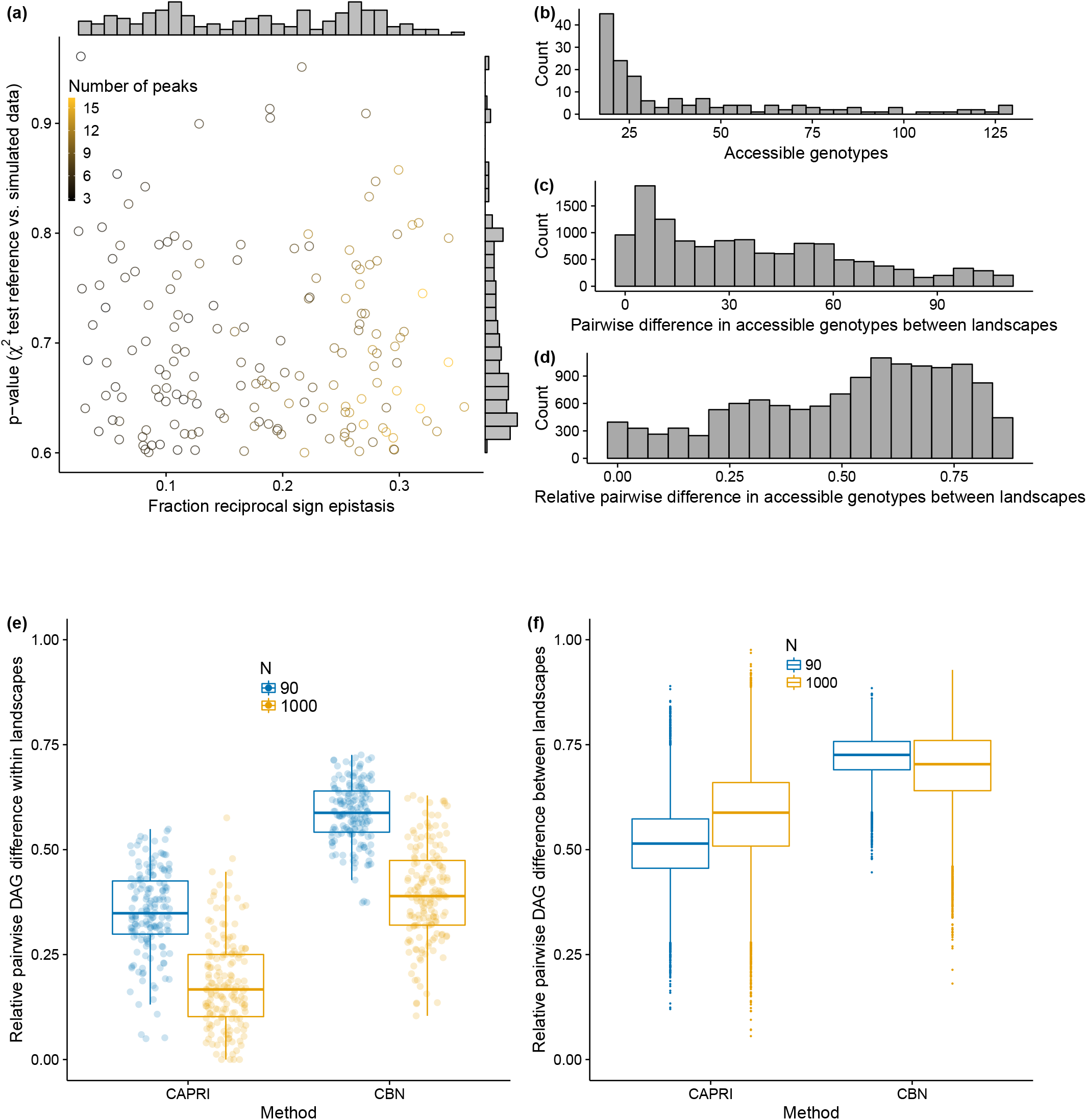
Fitness landscapes characteristics and DAG differences for pancreatic cancer. (a) p-value from the *χ*^2^ test comparing simulated vs. reference data set against reciprocal sign epistasis for the 162 fitness landscapes, with marginal histograms on the sides, and circles colored by number of peaks of the landscape. (b) Histogram of the number of accessible genotypes of the 162 fitness landscapes. (c) and (d): Histograms of pairwise and relative pairwise differences in accessible genotypes over all possible 162 * 161/2 pairs of landscapes. (e) Box-plots of relative pairwise differences over all pairs of the 20 DAGs inferred from each landscape with sample sizes of 90 (blue) and 1000 (orange) genotypes; thus, 162 points are shown, each the average of 20 * 19/2 pairwise differences. (f) Relative pairwise difference between all pairs of landscapes. Thus, each box-plot plot is based on 162 * 161/2 points; each point itself is the average of 400 (20 * 20) comparisons. (See Supplementary Material “Pancreas: results with other criteria” for these figures with different criteria).

As in section 2.3, there was considerable DAG-to-DAG variation within landscape, larger when using sample sizes of 90 genotypes instead of 1000 (Figure 5e). Much larger than the within-landscape DAG variation were the differences between DAGs inferred from different landscapes (Figure 5f): pairs of DAGs differed in a median of more than 50% of their edges, probably a consequence of the large differences between landscapes mentioned above (Figure 5c,d). In other words, this set of landscapes, all of which can produce genotype frequency data similar to the empirically observed one, not only showed large differences among themselves but lead to very different DAGs when evolutionary processes were run repeatedly on them.

## 3 Discussion

Cancer progression models (CPMs) assume restrictive fitness landscapes that, for instance, are devoid of reciprocal sign epistasis. Yet reciprocal sign epistasis may be common in cancer fitness landscapes [11]. What would be the consequences of using CPMs if tumors evolved on fitness landscapes that cannot be represented by DAGs from CPMs?

We saw (Figure 2) lower performance of cancer progression models in RMF landscapes relative to representable landscapes, and decreasing performance with increasing reciprocal sign epistasis in both RMF and DAG-derived landscapes. CPMs assume that acquiring a mutation in one gene does not decrease the probability of acquiring a mutation in another gene [34]. CBN, Oncogenetic Trees [15, 46], and CAPRESE [30] all share these features: even with unlimited data that faithfully represents the accessible genotypes in a landscape, they cannot fit non-representable fitness landscapes such as those that result from reciprocal sign epistasis. CAPRI can model XOR relationships [9, 40] and, thus, for example, synthetic lethality, but it requires specifying the XOR hypothesis *a priori*; therefore, it is not suitable for automated usage in non-representable landscapes. Separating patients into subtypes prior to analysis [9] cannot solve the problems caused by non-representable fitness landscapes because these problems are not the result of using CPMs on a mixture of individuals with different underlying fitness landscapes. This forces us to ask what is the meaning and how to interpret an inferred DAG in the presence of non-representability.

This paper also shows that there is a many-to-many relationship between DAGs inferred from CPMs and fitness landscapes, and this many-to-many relationship affects both representable and non-representable fitness landscapes. The consequences of this many-to-many relationship for the analysis of empirical data are illustrated with a pancreatic cancer data set. Genotype frequencies similar to the empirically observed one can be obtained under fitness landscapes that are very different from each other (Figure 5d) and that, when evolutionary processes are run on them, produce data that lead to the inference of DAGs that are also very different among themselves (Figure 5f). This many-to-many relationship between fitness landscapes and DAGs has two parts.

First, the same landscape generates data sets that often lead to inferring widely different DAGs (Figure 3). Even in representable landscapes with mutation and detection constant (Figure 3b, bottom row), DAGs differed in about 20% of their edges; this was using a sample size of N=1, 000 on landscapes of seven genes. More generally, we can think of a DAG as the output from applying a method on genotype frequency data that are the result of the function composition *Observed Genotype Frequencies = Detection ∘ Mutation ∘ Landscape*. The same landscape can lead to different observed genotype frequency data and, thus, different inferred DAGs if we change detection and mutation rates (see also [16]). When we did that (Figure 3b, top row), the DAGs inferred differed, on average, in 50% of their edges. Given that detection and mutation are likely to vary between tissues and cancer types [5, 25, 31], we could easily be inferring very different DAGs from similar underlying fitness landscapes.

Second, pairs of landscapes that differed in > 50% of their accessible genotypes (Figure 4) often lead to the same DAG, what we called “one DAG, many landscapes”. This is unavoidable if we use DAGs on non-representable landscapes: the number of fitness landscapes (as sets of accessible genotypes) is much larger than the number of possible cancer progression DAGs. And it probably becomes unavoidable too for representable landscapes when landscapes can lead to very different DAGs. So even under favourable conditions (representable landscapes, large sample sizes) the cross-sectional data from evolutionary processes is too variable and coarse to allow us to discern very different landscapes from each other (see also [43]). The results from the pancreatic cancer data probably exploit the ubiquitous many-to-many phenomenon, exacerbated by non-representability and sample sizes of N = 90. These results cast doubts on whether the restrictions inferred from any particular DAG capture those that exist in the underlying fitness landscape.

The many-to-many phenomenon affected both CAPRI and CBN, but the two methods were affected differently by mutation rate, detection regime and even landscape type (Figures 2 and 3). In particular, CAPRI’s DAG variability (Figure 3c,d) decreased in the RMF landscapes; this, however, does not mean that the quality of inferences is better in those landscapes, as it is not (see Figure 2b,c), but simply that is less variable around the error. The contrast in the behavior of CBN and CAPRI is probably caused by the different procedures that CAPRI and CBN use to accommodate noise and deviations from the fitted models. But there is an additional difference that affects interpretation. CBN (like Oncogenetic Trees [15, 46] and CAPRESE [30]) should only place an arrow between two genes, *A* and *C*, if *C* cannot be observed without *A* (*P*(*C*|*A*) > 0, *P*(*C*|¬*A*) = 0). This, however, is not necessarily the case for CAPRI (the condition *P*(*C*|*A*) > *P*(*C*|¬*A*) has to be true, but *P*(*C*|¬*A*) can be > 0). The arrows in CAPRI’s models, as explained in [40, p. 3017] “(…) imply ‘probability raising’ (…) [which] signifies that the presence of the earlier genomic alteration (…) that is advantageous in a Darwinian competition scenario increases the probability with which a subsequent advantageous genomic alteration (…) appears in the clonal evolution of the tumor.”. Therefore, with CAPRI a simple interpretation of arrows as “needed for occurrence” is precluded. The meaning of the arrows can depend on details of mutation rates and fitness differences between genotypes in ways that are difficult to reason about and that mix very different types of epistasis. For example, in the fitness landscape of Figure 1h, CAPRI could make *C* depend on *A* even if both were observed as single mutants (and even when the fitness of a mutant with only *C* is much larger than that of a mutant with only *A*) if the mutation rate of *C* was sufficiently small compared to *A*. Under identical mutation rates of *A* and *C*, CAPRI could make *A* depend on *C* even if both were observed as single mutants (e.g., if *C* were observed alone much more frequently than *A* because of clonal interference). Or, also under identical mutation rates, CAPRI, but also CBN, could even make *C* depend on *A* if a mutation in *C* lead to a very quick and large growth of the tumor that, because of the much larger population size, acquires an *A* mutation soon afterwards; here mutating *C* facilitates acquiring *A* not because of epistasis, but simply because *C*’s mutation increases population size. Making *C* depend on *A* would arguably be reverting causes and effects. As said above, CBN is not immune to these patterns. Moreover, in all of these cases the fitness of the double mutant *A*, *C* is larger than the fitness of each single mutant but there is no sign epistasis; in fact, the landscape of the figure is compatible both with magnitude epistasis of *A* and *C* and with no epistasis at all. This difference in the meaning of the arrows is probably the reason behind some of the counter-intuitive behavior of CAPRI and also explains CAPRI’s inference of many linear paths (see Supplementary Material “Plots of fitness landscapes and inferred DAGs”) and its larger PNDs. False negatives means failing to predict as possible an accessible genotype, and are the result of DAGs that encode too many restrictions, dependencies between genes that do not hold in the fitness landscapes. So, whereas we might try to map CBN’s output to structural features of the fitness landscape and to fundamentally different kinds of epistasis (magnitude, sign, reciprocal sign) this is not possible with CAPRI and emphasizes the different interpretation of CAPRI’s output even in representable fitness landscapes. More generally, it warns us that the meaning of the arrows can depend, in counter-intuitive ways, on details about the mutation rates, evolutionary dynamics, and their intersection with the detection process, details about which we know little; both CBN and CAPRI can be affected by these problems.

In conclusion, how much of a problem are non-representable fitness landscapes and the many-to-many phenomenon for using the inferences obtained by CPMs, for instance to identify therapeutic targets to block the progress of disease? Cancer progression models produce at best blurry maps of the underlying epistatic relationships in the fitness landscape, and differences in the underlying fitness landscape can have a large effect in the evolutionary dynamics of cancer and, thus, our opportunities for blocking the progress of disease. This raises the questions of whether we can asses from empirical data if landscapes are representable and, more importantly, what are the biomedical implications of errors and variability in the inferences of restrictions; even for representable landscapes, it might be extremely difficult to identify the correct dependency relationships between genes and, thus, the possible tumor progression paths, from cross-sectional data.

## 4 Methods

### 4.1 Generating random fitness landscapes

All DAGs and fitness landscapes in this paper use seven genes. That is the number of genes in the pancreatic cancer data set, the execution time of CBN increases steeply with number of genes beyond about seven genes [16], seven genes is probably close to the upper limits of fitness landscapes that can be easily visualized and related to their true DAGs (see Supplementary Material, “Plots of fitness landscapes and inferred DAGs”), and if number of genes has an effect on the problems reported in this paper they are likely to become worse with increasing numbers of genes.

The generation of the representable fitness landscapes had two steps (see also Supplementary Material, “Random fitness landscapes”): a) generating a random DAG of genes; b) assigning fitness values to mutations encoded in the DAG. To generate random DAGs, seven genes were first randomly split in 2 to 5 levels and each gene was randomly connected (as descendant) to randomly chosen genes (ancestors) from previous levels. The final DAG is the transitive reduction of the above generated DAG. From this DAG a fitness landscape was produced by: b.1) setting to 10^−9^ the fitness of any genotype that is not possible under the DAG (this makes it almost impossible to ever see a genotype of that kind —see supplementary material, “Paths through non-accessible genotypes”); b.2) assigning to every mutation with its dependencies satisfied (as specified by DAG) a random fitness effect uniformly distributed between 0.1 and 0.7 (these are values within those previously used in the literature [16, 33]). To give an example, suppose the DAG specifies that mutating gene *C* requires having genes A and B mutated, and genes A and B require no previous mutation; suppose fitness effects of mutations (with restrictions satisfied) are 0.1, 0.2, 0.3, respectively. The fitness of genotypes will be: A: 1.1 (=1 + 0.1); B: 1.2; AB: 1.1 * 1.2; C, AC, BC: 10^−9^ (since dependencies are not satisfied); ABC: 1.1 * 1.2 * 1.3, with wild-type always having fitness of 1. Note that, as shown by the example, converging or multiple arrows were given the meaning of an AND logical operator, not an OR.

Generation of DAG-derived but not representable fitness landscapes started by generating a representable DAG-derived fitness landscape as just described. Then, a randomly chosen subset of genotypes with two or more mutations and accessible under the DAG was made inaccessible. This fitness landscape was then checked to ensure that all seven genes were present in at least one accessible genotype.

The 200 RMF-derived non-representable fitness landscapes were obtained from a Rough Mount Fuji model [20, 35, 47] where the reference genotype and the decrease in fitness of a genotype per each unit increase in Hamming distance from the reference genotype were randomly chosen (see Supplementary Material, “Random fitness landscapes”). This gives a wide variety of fitness landscapes that encompass from close to additive to House of Cards models. Each fitness landscape was checked to ensure that all seven genes were present in at least one accessible genotype.

### 4.2 Evolutionary simulations, mutation rates, and detection regimes

I simulated evolution on the fitness landscapes using the model in [33], a logistic-like model where death rate depends on total population size. In addition to fitness (more precisely, birth rates) which are given by the fitness landscape, when using this model we need to specify initial population sizes, mutation rates, and detection/stopping conditions. Initial population size was set at 2,000 to ensure that most simulations reach cancer [33]; crossing fitness valleys with these initial sizes was extremely unlikely (see Supplementary Material “Paths through non-accessible genotypes”). Three different conditions of mutation rates were used: common mutation rate of rate of 1 × 10^−5^, common mutation rate of 1 × 10^−6^, and gene-specific mutation rates of 5.848 × 10^−6^, 5 × 10^−5^, 3.42 × 10^−6^, 2.924 × 10^−5^, 1.71 × 10^−5^, 2 × 10^−6^ and 1 × 10^−5^ for genes A to G, respectively; these mutation rates have a geometric mean of 1 × 10^−5^, with maximum spread between successive values within the maximum and minimum of 5 × 10^−5^ and 2 × 10^−6^, respectively. These are mutation rates within ranges previously used in the literature [7, 33], with a bias towards larger numbers (since we use only seven genes relevant for population growth). Each simulation was stopped when the tumor was detected and a whole tumor sample was taken. The probability of tumor detection increased with total tumor size and two conditions were used: the “fast” and the “slow” condition that correspond to detection probabilities of 0.1 and 0.01 when tumor size was 4.000 —twice the initial population size— (see Supplementary Material “Simulations: parameters and detection”).

### 4.3 Random fitness landscapes and simulations for the pancreatic cancer data

I repeatedly simulated data using a modified Rough Mount Fuji random fitness landscape model where the observed genotype combinations in the empirical data were guaranteed to be accessible. The procedure to create fitness landscapes involved two steps. First, fitness values were randomly assigned to the observed genotypes in the empirical data so that the observed genotypes were guaranteed to be accessible; this resulted in fitness values for the 18 observed genotypes. Second, and independently of the first step, a Rough Mount Fuji fitness landscape was generated (see 4.1 and supplementary material) but the fitnesses of the observed genotypes were set to those obtained in the first step. This guarantees that, regardless of the fitness of the rest of the genotypes in the landscape, the genotypes in the reference data set remain accessible. Generating fitness values so that the observed genotypes in the empirical data are accessible was done by creating a modified fitness graph [13] where each observed genotype was made the descendant of another observed genotype with one fewer mutation; iterating over genotypes, the fitness of each genotype was set to the sum of the genotype of its immediate ancestor plus a random uniform deviate. Once the fitness landscape had been generated, I simulated 90 genotypes from it (see section 4.2) using randomly selected parameters for mutation rates and detection regime (see Supplementary Material); I compared the genotype distributions with a chi-square test, keeping those where the p-value was > 0.6 (for both the reference *χ*^2^ distribution and a permutation test, because of possible cells of low counts) and all seven genes had been observed.

### 4.4 Measures of DAG performance and variability, landscape variability, and reciprocal sign epistasis

I used CBN [21, 22] and CAPRI [40] with their default settings (see Supplementary Material “Cancer progression methods and other sotware”) to obtain inferred DAGs. Reciprocal sign epistasis is the fraction of all pairs of mutations that had reciprocal sign epistasis (i.e., both mutations have an opposite effects on the other background) [8, 19] and was computed using MAGELLAN [8].

The **distance between two DAGs** is the number of the edges that differ between the transitive reduction of the two DAGs (the same as the sum of the absolute values of the entries in the matrix difference of the adjacency matrices of two DAGs or, equivalently, the cardinality of the symmetric difference of the sets of edges of the DAGs). CAPRI can return DAGs that contain both direct and indirect edges between nodes (i.e., DAGs that contain more edges than a similar smaller DAG with the same reachability conditions): all comparisons involved the transitive reduction of the DAGs. The **relative distance between DAGs**, used to assess DAG-to-DAG variability, is the distance between two DAGs divided by the total number of distinct edges in the two DAGs (i.e., the cardinality of the union of the sets of edges of the two DAGs); this provides an easy to interpret measure in the range [0,1] where 0 means identical DAGs and 1 means all edges are different. **PND** genotype mispredictions is the ratio of false negative genotype mispredictions over the total number of genotypes that are accessible in a landscape. A genotype, even if accessible under a given landscape, might actually never be observable under small mutation rates and fast detection regimes. To avoid penalizing inferences for non-observable genotypes, I corrected the count of false negatives using, as reference, not the full set of accessible genotypes under a fitness landscape, but only the subset of accessible genotypes that had been observed with a frequency larger than 5 in 1000 (measured on the 20,000 simulations) so that the probability of not observing the minimal frequency genotype in a sample of 1,000 genotypes is less than 1%. PND is equivalent to 1 – recall or 1 – sensitivity. **PFD** genotype mispredictions is the ratio of false positive genotype mispredictions over the total number of genotypes that can exist according to the DAG; this is equivalent to 1 – precision or 1 – positive predictive value.

The **pairwise difference of accessible genotypes** between two landscapes is the sum of the number of genotypes accessible under one landscape and not accessible under the other (i.e., the cardinality of the symmetric difference, or disjunctive union, of the sets of accessible genotypes of each landscape). The **relative pairwise difference of accessible genotypes** divides the pairwise difference by the total number of distinct accessible genotypes in the two landscapes (i.e., the cardinality of the of the union of the sets of accessible landscapes of the two landscapes).

### 4.5 Linear mixed-effects models

For the analysis of the simulations on the 500 fitness landscapes, I used linear mixed-effects models to examine how type of landscape, mutation rate, detection scheme, amount of reciprocal sign epistasis, and number of accessible genotypes, affected PND, PFD, and the two relative pairwise DAG distances (Supplementary Material “Linear mixed-effects models”). Models used fitness landscape as random effect except for relative pairwise DAG distance over mutation and detection as here, since I averaged over all mutation and detection regimes, only a single value per landscape is used. Type of landscape, number of accessible genotypes, and amount of reciprocal sign epistasis are always the same for a landscape.

For ease of interpretation (i.e., to avoid presenting coefficients from models with three and four way interactions), Figures 2 and 3 present results from models for each method and landscape type separately, but I also fitted a single model to assess the interactions of method and landscape type with the other factors. To assess the effects of method and type of landscape independently without considering any other factors (i.e., to examine the significance of the differences in Figures 2a,b and Figure 3b), I fitted models than only had landscape type and method as explanatory variables (these models are of course equivalent to multistratum models for split-plot designs [17, 37]). In all cases, models were fitted using sum-to-zero contrasts: each main effect parameter is to be interpreted as the (marginal) deviation of that level from the overall mean, and the interaction parameter as the deviation of the linear predictor of the cell mean (for that combination of levels) from the addition of the corresponding main effect parameters (see [32]). Continuous regressors (reciprocal sign epistasis and number of accessible genotypes) were scaled (mean 0, variance 1) to allow for a simpler interpretation of the magnitude of their coefficients; the intercept term is to be interpreted as the predicted response at the average value of the regressors.

## 5 Acknowledgments

C. Lazaro-Perea and I.B. Lightwood for comments on the ms. S. Alvarez-Tolcheff and C. Vasallo for discussion and comments about the ms. J. Poyatos for discussion and suggestions about figures. G. Caravagna for discussion about progression models and help understanding CAPRI.

## References

[1] Beerenwinkel, N., Antal, T., Dingli, D., Traulsen, A., Kinzler, K. W., Velculescu, V. E., Vogelstein, B., Nowak, M. A., 2007. Genetic progression and the waiting time to cancer. PLoS computational biology, 3(11):e225. doi:10.1371/journal.pcbi.0030225. URL http://www.pubmedcentral.nih.gov/articlerender.fcgi?artid=2065895&tool=pmcentrez&rendertype=abstract.

[2] Beerenwinkel, N., Greenman, C. D., Lagergren, J., 2016. Computational Cancer Biology: An Evolutionary Perspective. PLoS Comput. Biol., 12(2):e1004717. doi:10.1371/journal.pcbi.1004717.

[3] Beerenwinkel, N., Schwarz, R. F., Gerstung, M., Markowetz, F., 2014. Cancer evolution: Mathematical models and computational inference. Systematic Biology, 64(1):e1–e25. doi: 10.1093/sysbio/syu081. URL http://sysbio.oxfordjournals.org/content/64/1/e1.

[4] Beijersbergen, R. L., Wessels, L. F. A., Bernards, R., 2017. Synthetic Lethality in Cancer Therapeutics. Annual Review of Cancer Biology, 1(1):141–161. doi:10.1146/annurev-cancerbio-042016-073434. URL http://dx.doi.org/10.1146/annurev-cancerbio-042016-073434.

[5] Blokzijl, F., De Ligt, J., Jager, M., Sasselli, V., Roerink, S., Sasaki, N., Huch, M., Boymans, S., Kuijk, E., Prins, P., Nijman, I. J., Martincorena, I., Mokry, M., Wiegerinck, C. L., Middendorp, S., Sato, T., Schwank, G., Nieuwenhuis, E. E. S., Verstegen, M. M. A., van der Laan, L. J. W., de Jonge, J., IJzermans, J. N. M., Vries, R. G., van de Wetering, M., Stratton, M. R., Clevers, H., Cuppen, E., van Boxtel, R., 2016. Tissue-specific mutation accumulation in human adult stem cells during life. Nature, 538(7624) :260–264. doi: 10.1038/nature19768.

[6] Blomen, V. A., Májek, P., Jae, L. T., Bigenzahn, J. W., Nieuwenhuis, J., Staring, J., Sacco, R., van Diemen, F. R., Olk, N., Stukalov, A., Marceau, C., Janssen, H., Carette, J. E., Bennett, K. L., Colinge, J., Superti-Furga, G., Brummelkamp, T. R., 2015. Gene essentiality and synthetic lethality in haploid human cells. Science, 350(6264) :1092–1096. doi:10.1126/science.aac7557. URL http://www.sciencemag.org/content/350/6264/1092.

[7] Bozic, I., Antal, T., Ohtsuki, H., Carter, H., Kim, D., Chen, S., Karchin, R., Kinzler, K. W., Vogelstein, B., Nowak, M. A., 2010. Accumulation of driver and passenger mutations during tumor progression. Proceedings of the National Academy of Sciences of the United States of America, 107:18545–18550. doi:10.1073/pnas.1010978107. URL http://www.ncbi.nlm.nih.gov/pubmed/20876136.

[8] Brouillet, S., Annoni, H., Ferretti, L., Achaz, G., 2015. MAGELLAN: A tool to explore small fitness landscapes. bioRxiv, p. 031583. doi:10.1101/031583. URL http://biorxiv.org/content/early/2015/11/13/031583.

[9] Caravagna, G., Graudenzi, A., Ramazzotti, D., Sanz-Pamplona, R., Sano, L. D., Mauri, G., Moreno, V., Antoniotti, M., Mishra, B., 2016. Algorithmic methods to infer the evolutionary trajectories in cancer progression. PNAS, p. 201520213. doi:10.1073/pnas.1520213113. URL http://www.pnas.org/content/early/2016/06/27/1520213113.

[10] Cheng, Y.-K., Beroukhim, R., Levine, R. L., Mellinghoff, I. K., Michor, F., 2011. Reply to Parsons: Many tumor types follow the monoclonal model of tumor initiation. Proceedings of the National Academy of Sciences, 108(5):E16–E16. doi:10.1073/pnas.1018584108. URL http://www.pnas.org/cgi/doi/10.1073/pnas.1018584108.

[11] Chiotti, K. E., Kvitek, D. J., Schmidt, K. H., Koniges, G., Schwartz, K., Donckels, E. A., Rosenzweig, F., Sherlock, G., 2014. The Valley-of-Death: Reciprocal sign epistasis constrains adaptive trajectories in a constant, nutrient limiting environment. Genomics, 104(6, Part A):431–437. doi:10.1016/j.ygeno.2014.10.011. URL http://www.sciencedirect.com/science/article/pii/S0888754314002122.

[12] Cristea, S., Kuipers, J., Beerenwinkel, N., 2016. pathTiMEx: Joint Inference of Mutually Exclusive Cancer Pathways and Their Progression Dynamics. Journal of Computational Biology. doi:10.1089/cmb.2016.0171. URL http://online.liebertpub.com/doi/abs/10.1089/cmb.2016.0171.

[13] Crona, K., Greene, D., Barlow, M., 2013. The peaks and geometry of fitness landscapes. Journal of Theoretical Biology, 317:1–10. doi:10.1016/j.jtbi.2012.09.028. URL http://www.sciencedirect.com/science/article/pii/S0022519312005061.

[14] de Visser, J. A. G. M., Krug, J., 2014. Empirical fitness landscapes and the predictability of evolution. Nat Rev Genet, 15(7):480–490. doi:10.1038/nrg3744. URL http://www.nature.com/nrg/journal/v15/n7/full/nrg3744.html.

[15] Desper, R., Jiang, F., Kallioniemi, O. P., Moch, H., Papadimitriou, C. H., Schäffer, A. A., 1999. Inferring tree models for oncogenesis from comparative genome hybridization data. J Comput Biol, 6(1):37–51. URL http://view.ncbi.nlm.nih.gov/pubmed/10223663.

[16] Diaz-Uriarte, R., 2015. Identifying restrictions in the order of accumulation of mutations during tumor progression: Effects of passengers, evolutionary models, and sampling. BMC Bioinformatics, 16(41):0–36. doi:doi:10.1186/s12859–015–0466–7. URL http://www.biomedcentral.com/1471–2105/16/41/abstract.

[17] Faraway, J. J., 2016. Extending the Linear Model with R: Generalized Linear, Mixed Effects and Nonparametric Regression Models, Second Edition. Chapman and Hall/CRC, Boca Raton, 2 edition edition. ISBN 978-1-4987-2096-0.

[18] Fearon, E., Vogelstein, B., 1990. A genetic model for colorectal tumorigenesis. Cell, 61:759–767. URL http://www.sciencedirect.com/science/article/pii/009286749090186I.

[19] Ferretti, L., Schmiegelt, B., Weinreich, D., Yamauchi, A., Kobayashi, Y., Tajima, F., Achaz, G., 2016. Measuring epistasis in fitness landscapes: The correlation of fitness effects of mutations. Journal of Theoretical Biology, 396:132–143. doi:10.1016/j.jtbi.2016.01.037. URL http://www.sciencedirect.com/science/article/pii/S0022519316000771.

[20] Franke, J., Klözer, A., de Visser, J. A. G. M., Krug, J., 2011. Evolutionary Accessibility of Mutational Pathways. PLoS Comput Biol, 7(8):e1002134. doi:10.1371/journal.pcbi.1002134. URL http://dx.doi.org/10.1371/journal.pcbi.1002134.

[21] Gerstung, M., Baudis, M., Moch, H., Beerenwinkel, N., 2009. Quantifying cancer progression with conjunctive Bayesian networks. Bioinformatics (Oxford, England), 25(21):2809–2815. doi:10.1093/bioinformatics/btp505. URL http://dx.doi.org/10.1093/bioinformatics/btp505%0020http://www.bsse.ethz.ch/cbg/software/ct-cbn.

[22] Gerstung, M., Eriksson, N., Lin, J., Vogelstein, B., Beerenwinkel, N., 2011. The Temporal Order of Genetic and Pathway Alterations in Tumorigenesis. PLoS ONE, 6(11):e27136. doi:10.1371/journal.pone.0027136. URL http://dx.plos.org/10.1371/journal.pone.0027136%0020http://www.bsse.ethz.ch/cbg/software/ct-cbn.

[23] Greaves, M., 2015. Evolutionary Determinants of Cancer. Cancer Discovery, 5(8):806–820. doi:10.1158/2159-8290.CD-15-0439. URL http://cancerdiscovery.aacrjournals.org/content/5/8/806.

[24] Hainke, K., Rahnenführer, J., Fried, R., 2012. Cumulative disease progression models for cross-sectional data: A review and comparison. Biometrical journal. Biometrische Zeitschrift, 54(5):617–40. doi:10.1002/bimj.201100186. URL http://www.ncbi.nlm.nih.gov/pubmed/22886685.

[25] Hao, D., Wang, L., Di, L.-j., 2016. Distinct mutation accumulation rates among tissues determine the variation in cancer risk. Scientific Reports, 6:19458. doi:10.1038/srep19458. URL http://www.nature.com/srep/2016/160120/srep19458/full/srep19458.html.

[26] Jones, S., Zhang, X., Parsons, D. W., Lin, J. C.-H., Leary, R. J., Angenendt, P., Mankoo, P., Carter, H., Kamiyama, H., Jimeno, A., Hong, S.-M., Fu, B., Lin, M.-T., Calhoun, E. S., Kamiyama, M., Walter, K., Nikolskaya, T., Nikolsky, Y., Hartigan, J., Smith, D. R., Hidalgo, M., Leach, S. D., Klein, A. P., Jaffee, E. M., Goggins, M., Maitra, A., Iacobuzio-Donahue, C., Eshleman, J. R., Kern, S. E., Hruban, R. H., Karchin, R., Papadopoulos, N., Parmigiani, G., Vogelstein, B., Velculescu, V. E., Kinzler, K. W., 2008. Core signaling pathways in human pancreatic cancers revealed by global genomic analyses. Science (New York, N.Y.), 321(5897):1801–6. doi:10.1126/science.1164368. URL http://www.ncbi.nlm.nih.gov/pubmed/18772397.

[27] Krzywinski, M., Schein, J., Birol, b., Connors, J., Gascoyne, R., Horsman, D., Jones, S. J., Marra, M. A., 2009. Circos: An information aesthetic for comparative genomics. Genome Research, 19(9):1639–1645. doi:10.1101/gr.092759.109. URL http://dx.doi.org/10.1101/gr.092759.109.

[28] Leung, A. W. Y., de Silva, T., Bally, M. B., Lockwood, W. W., 2016. Synthetic lethality in lung cancer and translation to clinical therapies. Molecular Cancer, 15:61. doi:10.1186/s12943-016-0546-y. URL http://dx.doi.org/10.1186/s12943-016-0546-y.

[29] Lipinski, K. A., Barber, L. J., Davies, M. N., Ashenden, M., Sottoriva, A., Gerlinger, M., 2016. Cancer Evolution and the Limits of Predictability in Precision Cancer Medicine. Trends in Cancer, 2(1):49–63. doi:10.1016/j.trecan.2015.11.003. URL http://linkinghub.elsevier.com/retrieve/pii/S2405803315000692.

[30] Loohuis, L. O., Caravagna, G., Graudenzi, A., Ramazzotti, D., Mauri, G., Antoniotti, M., Mishra, B., 2014. Inferring Tree Causal Models of Cancer Progression with Probability Raising. PLoS ONE, 9(10):e108358. doi:10.1371/journal.pone.0108358. URL http://dx.plos.org/10.1371/journal.pone.0108358%0020http://bimib.disco.unimib.it/index.php/Tronco%0020http://journals.plos.org/plosone/article?id=10.1371/journal.pone.0108358.

[31] Martincorena, I., Campbell, P. J., 2015. Somatic mutation in cancer and normal cells. Science, 349(6255):1483–1489. doi:10.1126/science.aab4082. URL http://www.sciencemag.org/content/349/6255/1483.

[32] McCullagh, P., Nelder, J., 1989. Generalized Linear Models, 2nd Ed. Chapman and Hall/CRC, London.

[33] McFarland, C. D., Korolev, K. S., Kryukov, G. V., Sunyaev, S. R., Mirny, L. A., 2013. Impact of deleterious passenger mutations on cancer progression. Proceedings of the National Academy of Sciences of the United States of America, 110(8):2910–5. doi: 10.1073/pnas.1213968110. URL http://www.ncbi.nlm.nih.gov/pubmed/23388632.

[34] Misra, N., Szczurek, E., Vingron, M., 2014. Inferring the paths of somatic evolution in cancer. Bioinformatics (Oxford, England), 30(17):2456–2463. doi:10.1093/bioinformatics/btu319. URL http://www.ncbi.nlm.nih.gov/pubmed/24812340.

[35] Neidhart, J., Szendro, I. G., Krug, J., 2014. Adaptation in Tunably Rugged Fitness Landscapes: The Rough Mount Fuji Model. Genetics, 198(2):699–721. doi:10.1534/genetics.114.167668. URL http://www.genetics.org/content/198/2/699.

[36] Parsons, B. L., 2011. Monoclonal tumor origin is an underlying misconception of the RESIC approach. Proceedings of the National Academy of Sciences of the United States of America, 108(5):E15; author reply E16. doi:10.1073/pnas.1017998108. URL http://www.pubmedcentral.nih.gov/articlerender.fcgi?artid=3033296&tool=pmcentrez&rendertype=abstract.

[37] Pinheiro, J. C., Bates, D. M., 2000. Mixed-Effects Models in S and S-PLUS. Springer, New York. ISBN 978-0-387-98957-0 978-1-4419-0317-4.

[38] Poelwijk, F. J., Kiviet, D. J., Weinreich, D. M., Tans, S. J., 2007. Empirical fitness landscapes reveal accessible evolutionary paths. Nature, 445(7126):383–6. doi: http://dx.doi.org/10.1038/nature05451. URL http://search.proquest.com/docview/204523563/abstract/CED5FE881C9942BCPQ/1.

[39] Poelwijk, F. J., Tănase-Nicola, S., Kiviet, D. J., Tans, S. J., 2011. Reciprocal sign epistasis is a necessary condition for multi-peaked fitness landscapes. Journal of Theoretical Biology, 272(1):141–144. doi:10.1016/j.jtbi.2010.12.015. URL http://www.sciencedirect.com/science/article/pii/S0022519310006703.

[40] Ramazzotti, D., Caravagna, G., OlDe Loohuis, L., Graudenzi, A., Korsunsky, I., Mauri, G., Antoniotti, M., Mishra, B., 2015. CAPRI: Efficient inference of cancer progression models from cross-sectional data. Bioinformatics, 31(18):3016–3026. doi:10.1093/bioinformatics/btv296. URL https://academic.oup.com/bioinformatics/article-lookup/doi/10.1093/bioinformatics/btv296.

[41] Sniegowski, P. D., Gerrish, P. J., 2010. Beneficial mutations and the dynamics of adaptation in asexual populations. Philosophical Transactions of the Royal Society of London B: Biological Sciences, 365(1544):1255–1263. doi:10.1098/rstb.2009.0290. URL http://rstb.royalsocietypublishing.org/content/365/1544/1255.

[42] Sprouffske, K., Merlo, L. M. F., Gerrish, P. J., Maley, C. C., Sniegowski, P. D., 2012. Cancer in light of experimental evolution. Current biology : CB, 22(17):R762–71. doi: 10.1016/j.cub.2012.06.065. URL http://www.pubmedcentral.nih.gov/articlerender.fcgi?artid=3457634&tool=pmcentrez&rendertype=abstract.

[43] Sprouffske, K., Pepper, J. W., Maley, C. C., 2011. Accurate reconstruction of the temporal order of mutations in neoplastic progression. Cancer prevention research (Philadelphiaa, Pa.), 4(7):1135–44. doi:10.1158/1940-6207.CAPR-10–0374. URL http://www.pubmedcentral.nih.gov/articlerender.fcgi?artid=3131446&tool=pmcentrez&rendertype=abstract.

[44] Srivas, R., Shen, J. P., Yang, C. C., Sun, S. M., Li, J., Gross, A. M., Jensen, J., Licon, K., Bojorquez-Gomez, A., Klepper, K., Huang, J., Pekin, D., Xu, J. L., Yeerna, H., Sivaganesh, V., Kollenstart, L., van Attikum, H., Aza-Blanc, P., Sobol, R. W., Ideker, T., 2016. A Network of Conserved Synthetic Lethal Interactions for Exploration of Precision Cancer Therapy. Molecular Cell, 63(3):514–525. doi:10.1016/j.molcel.2016.06.022. URL http://www.cell.com/molecular-cell/abstract/S1097-2765(16)30280-5.

[45] Stephens, P. J., Greenman, C. D., Fu, B., Yang, F., Bignell, G. R., Mudie, L. J., Pleasance, E. D., Lau, K. W., Beare, D., a. Stebbings, L., 2011. Massive Genomic Rearrangement Acquired in a Single Catastrophic Event during Cancer Development. Cell, 144(1):27–40. doi:10.1016/j.cell.2010.11.055. URL http://linkinghub.elsevier.com/retrieve/pii/S0092867410013772.

[46] Szabo, A., Boucher, K. M., 2008. Oncogenetic trees. In Tan, W.-Y., Hanin, L., editors, Handbook of Cancer Models with Applications, pp. 1–24. World Scientific. URL http://www.worldscibooks.com/lifesci/6677.html.

[47] Szendro, I. G., Franke, J., de Visser, J. A. G. M., Krug, J., 2013. Predictability of evolution depends nonmonotonically on population size. PNAS, 110(2):571–576. doi:10.1073/pnas.1213613110. URL http://www.pnas.org/content/110/2/571.

[48] Weinreich, D. M., Watson, R. A., Chao, L., 2005. Perspective: Sign epistasis and genetic constraint on evolutionary trajectories. Evolution, 59(6):1165–1174.

